# Water quality and microbial load: a double-threshold identification procedure intended for space applications

**DOI:** 10.1101/377697

**Authors:** Stefano Amalfitano, Caterina Levantesi, Laurent Garrelly, Donatella Giacosa, Francesca Bersani, Simona Rossetti

## Abstract

During longer-lasting future space missions, water renewal by ground-loaded supplies will become increasingly expensive and unmanageable for months. Space exploration by self-sufficient space-crafts is thus demanding the development of culture-independent microbiological methods for in-flight water monitoring to counteract possible contamination risks. In this study, we aimed at evaluating microbial load data assessed by selected techniques with current or promising perspectives in space applications (i.e., HPC, ATP-metry, qPCR, flow cytometry), through the analysis of water sources with constitutively different contamination levels (i.e., chlorinated and unchlorinated tap waters, groundwaters, river waters, wastewaters). Using a data-driven double-threshold identification procedure, we identified and presented new alternative standards of water quality based on the assessment of the total microbial load. Our approach is suitable to provide an immediate alert of microbial load peaks, thus enhancing the crew responsiveness in case of unexpected events due to water contamination and treatment failure. Finally, the backbone dataset could help in managing water quality and monitoring issues for both space and Earth-based applications.

## 1 Introduction

Aquatic microbes are retained as primary constituents of all known water sources aboard the International Space Station (ISS), as well as in future human spaceflights and planetary outposts (Horneck et al., 2010). Since space missions are expected to become longer lasting, space exploration is demanding the development of methods for in-flight monitoring, suitable to face microbial contamination risks within human confined conditions (Allen et al., 2018; Karouia et al., 2017; Yamaguchi et al., 2014). NASA has been developed microbial control strategies to minimize detrimental cell growth during spaceflight by reducing humidity, eliminating free water, and maintaining high-volume exchange and air filtration. The ISS is maintained at around 22°C with a relative humidity of around 60%, with pressure and oxygen concentrations very close to those at sea level on Earth (Pierson et al., 2013).

Currently on ISS, water samples are archived every 6 months for further post-flight analysis. In addition, samples are processed in-flight once every three months with the US-supplied Water Microbiology Kit for the quantification of heterotrophic bacteria (Heterotrophic Plate Counts - HPC) and the presence of coliforms (Van Houdt et al., 2012). Leaving aside the evidence that the microbial biomass is mainly composed by viable but not cultivable microorganisms (Colwell, 2009), the development of culture-independent methods for space applications is pushed fundamentally by the requirement of timeliness of results and the need to avoid microbial regrowth from analytical wastes. Candidate methods have to comply with limitations in volume, working time, power, safety and microgravity, thus being suitable for automation, lightweight and with minimal consumables. Since first experiments conducted in space, bioluminescence and PCR-based methods have been tested for monitoring the microbial load under microgravity conditions (Castro et al., 2004; Guarnieri et al., 1997; La Duc et al., 2004). Currently, joint scientific and industrial efforts have been focused on developing an on-line self-loading ATP-based monitoring module within an integrated breadboard system to control microbial contamination in water systems during human spaceflights. The module contains ATP-releasing reagents to lyse cells and release ATP, which reacts with d-luciferin in presence of luciferase to produce detectable light signals. The light intensity is then measured as Relative Light Units (RLU), which can be interpreted as a measure of ATP concentration (i.e., H2020 EU project “Biocontamination Integrated Control of Wet Systems for Space Exploration”, http://biowyse.eu/). Moreover on the ISS, biomolecular methods and sample processing for DNA extraction and gene sequencing has been tested within dedicated projects (e.g., Genes in Space-3, Wet-lab2) and through customized devices (e.g., miniPCR, MinION, Razor EX PCR) (Boguraev et al., 2017; Karouia et al., 2017; Parra et al., 2017). The quantitative real-time PCR (qPCR) could be applied to assess the absolute bacterial abundance by measuring the number of 16S rRNA housekeeping gene copies in the total DNA extracted from a water sample (Smith and Osborn, 2009). Among the consolidated water monitoring approaches for the direct quantification of aquatic microorganisms, flow cytometry (FCM) has to be also considered, since it was defined as an unparalleled high-throughput technology for single cell counting and characterization in a panoply of applications (Robinson and Roederer, 2015). The basic cytometric detection combines laser light scatter and fluorescence signals, with the ability to discriminate microbial cell subpopulations, phenotypes (e.g., size and shape), and constitutive properties detected upon specific staining procedures (e.g., per-cell nucleic acid content) (Wang et al., 2010).

By largely disregarding recent monitoring techniques and their methodological improvements, current standards for microbiological evaluations are set on the occurrence of few microorganisms, indicators of fecal pollution and hence of the possible co-presence of pathogenic species (European Union, 1998). Microbial specifications and monitoring requirements for ISS waters have fixed the limit of HPC ≤50 CFU/ml to meet the onboard quality standards in U.S. and Russian segments (Duncan et al., 2008). However, the space water microbiology was recently pushed beyond the standardized cultivation-based methods (Moissl-eichinger et al., 2016), also due to the finding that spaceflight microgravity conditions provided conflicting results, with insufficient and largely unpredictable indications on the microbial growth patterns and the virulence of opportunistic human pathogens (Huang et al., 2018).

In this study, we explored whether alternative methods to assess the water microbial load could be supportive of routine monitoring practices, thus challenging conventional heterotrophic plate counts in space applications. Through the analysis of water sources with constitutively different microbial loads (i.e., chlorinated and unchlorinated tap waters, groundwaters, river waters, wastewaters), we aimed to (i) cross-validate candidate techniques suitable to assess the onboard water microbial load and selected among those consolidated in terrestrial applications (i.e., HPC, ATP-metry, qPCR, FCM), and (ii) propose a data-driven procedure to determine new water quality standards based on the cultivation-independent assessment of the total microbial load.

By considering the high costs and logistic limitations of water renewal with Earth-supplied resources, we hypothesized that a double-threshold identification procedure could be applicable to identify water microbial contamination events and help counteracting health-risks, which could unexpectedly occur during longer-lasting space missions.

## 2 Material and methods

### 2.1 Selection and collection of water samples

Water samples were collected in one-liter sterile plastic bottles containing pre-dosed sodium thiosulfate (1 ml of 10% solution per bottle), transported in refrigerated boxes and stored at 5 ± 3°C for maximum 24 hours before analysis. A total of 35 samples was drawn from five types of water sources with a naturally different microbial load. Each water type was represented by seven independent samples: chlorinated Tap Water (cTW1-7), unchlorinated Tap Water (uTW1-7), Ground Water (GW1-7), River Water (RW1-7), and Waste Water (WW1-7).

Turbidity was measured in all samples and expressed as nephelometric turbidity units (NTU). The chlorinated tap waters (cTWs = 0.2-0.3 NTU) were collected along the drinking water distribution network, namely at the inlet of water kiosks in seven towns supplied by SMAT in the province of Turin (Italy). All cTWs were disinfected with sodium hypochlorite (final concentration 0.1-0.2 mg/l). Unchlorinated tap waters (uTWs = 0.1-0.5 NTU) were gathered at the granular activated carbon filter outlets of a drinking water treatment plant, following sedimentation, break-point chlorination, chlorine-dioxide peroxidation and clariflocculation with polyhydroxy aluminium chloride. Groundwater samples (GWs = 0.2-7.2 NTU) were collected from wells located in the urban area of Turin (Italy), before any kind of subsequent treatment. River water samples (RW = 0.2-1.8 NTU) were collected from water catchment areas upstream different drinking water treatment plants. Wastewater samples (WWs = 1.0-9.6 NTU) were collected from secondary effluents of activated-sludge wastewater treatment plants.

### 2.2 Plate cultivation and heterotrophic plate counts

Heterotrophic plate counts (HPC) were performed on 90-mm Petri dishes filled with either Yeast Extract Agar (YEA) medium (Sifin Diagnostics, Germany) or Reasoner’s 2A Agar (R2A) medium (Thermofisher Diagnostics, US) (APHA, 2005; ISO6222, 1999). Following lab incubation at 22°C for three or seven days, all results derived from the average of two plates, each within the countable range. Depending on the expected microbial concentrations, different volumes (from 1 ml to 100 ml) were analyzed in order to reach a measurable range in terms of colony forming units (CFUs). Volumes up to 1 ml were included in molten medium, while larger volumes (up to 100 ml) were filtered onto cellulose nitrate membranes (0.45-µm pore size; Millipore).

### 2.3 ATP-metry

The total intracellular ATP content was measured to estimate the microbial load using the technology developed by GL Biocontrol (Clapiers, France). Briefly, 2 drops of DENDRIAG reagent were added to the water sample (10-50 ml) and measured using the GL Biocontrol instrument to obtain the R1 (RLU) result. Then four drops of DENDRIAG reagent were dispensed into the plastic packaging of the filter and the reactive was backflush by pressing air through the filter. The reagent was pushed into the measuring tube and measured to obtain the R1 (RLU) result. Then, one drop of STANDARD 1000 reagent was added and measured to obtain the R2 (RLU) result. The concentration of intracellular ATP is given in picograms per milliliter, by the following calculations:

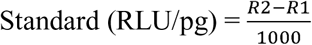

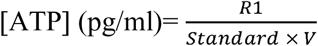

With R1 (RLU): sample result; R2 (RLU): sample + standard result; V (ml) volume of water analyzed.

### 2.4 DNA extraction and quantitative real-time PCR

The qPCR Sybr Green assay was utilized to measure the 16S rDNA gene copy number in 25 µl of sample using the CFX96 Touch Real Time PCR Detection System (Bio-Rad, USA). Reactions contained 5 µl of DNA template (from 50 to 5 ng DNA for reaction tube), 12.5 µL of 2X SYBR Green Supermix (Bio-Rad USA), and primers at required concentrations (Di Cesare et al., 2015). Triplicates samples and no template controls (NTCs) were analyzed. The *E.coli* 16S rDNA was used as positive control, and standard curves were produced with gene copy numbers from 10^2^ to 10^6^ genes per reaction tube. The concentration of the amplified DNA was determined using NanoDrop spectrophotometer. The gene copy number per µl of solution was calculated according to literature reports (Czekalski et al., 2012). The qPCR results were reported as the mean of measurements of triplicates analysis with standard deviations. Data were analyzed with the CFX ManagerTM software v3.1 (Bio-Rad, Italy).

### 2.5 Flow cytometry

The aquatic microbial cells were characterized by using the Flow Cytometer A50-micro (Apogee Flow System, Hertfordshire, England) equipped with a solid state laser set at 20 mV and tuned to an excitation wave length of 488 nm. The volumetric absolute counting was carried out on fixed (2% formaldehyde, final concentration) and unfixed water samples, stained with SYBR Green I (1:10000 dilution; Molecular Probes, Invitrogen) or with SYBR Green I and propidium iodide (PI = 10 µg ml^-1^, f.c.) for 10 min in the dark at room temperature. The light scattering signals (forward and side scatters), the green fluorescence (530/30 nm) and red fluorescence (> 610 nm) were acquired for the single cell characterization. A fluorescence threshold was set at 10 units on the green channel. Samples were run at low flow rates to keep the number of events below 1000 events per second. The total number of prokaryotic cells (i.e., total cell counts – TCC) was determined by their signatures in a plot of the side scatter vs the green fluorescence (Gasol and Morán, 2015). Live and dead cells were differentiated in a plot of green vs red fluorescence. Viable cells (i.e., intact cell counts - ICC) showed higher green fluorescence signals than the membrane compromised dead cells selectively marked in red by propidium iodide (Grégori et al., 2018). The instrumental settings were kept the same for all samples in order to achieve comparable data. The data were analyzed using the Apogee Histogram Software v2.05.

Total cell counts were double-checked by epifluorescence microscopy on all samples by following consolidated literature procedures (Porter and Feig, 1980). Briefly, aliquots of fixed samples were filtered through 0.2 μm polycarbonate filters (Ø 25 mm, Millipore) by gentle vacuum (<0.2 bar), and stained for 5 min with DAPI (4′, 6-diamidino-2-phenylindole; 1.5 µg ml^-1^ final concentration). Filters were stored at −20°C until microscope inspection. Total cell counts were performed by the epifluorescence microscope BX51 (Olympus, Germany) at 1500X magnification by counting a minimum of 300 cells in >10 microscopic fields randomly selected across each filter.

### 2.6 Data elaboration and statistical analysis

All data were log(x+1) transformed to facilitate comparability among parameters derived from different methods and waters sources. The nonparametric Kruskal-Wallis test, with Mann-Whitney post-hoc pairwise comparisons, was used to verify whether statistical differences in median values occurred among water groups according to each single parameter. The one-way nonparametric multivariate analysis of variance (PERMANOVA), based on the Euclidean distance measure, was used to test the overall significance of difference between water groups.

A frequency distribution model (FDM) of log-transformed data was applied to discriminate between two water groups, hereafter named waters with low and high microbial load. The number of bins was set at 4 for each single-parameter data series (Wand, 1997). Following the consolidated approach applied to assess upper confidence limits for natural background chemical concentrations (USEPA, 2002), a first confidence threshold (hereafter named warning threshold) was set as the 95th percentile of values assessed from the low microbial load group. A second and higher confidence threshold (hereafter named alarm threshold) was arbitrarily set as the 5^th^ percentile of values assessed from the high microbial load group.

The linear regression model (LRM) was applied in order to cross-validate the FDM thresholds using the possible combinations of log-transformed independent parameters (i.e., HPC-R2A vs ATP; HPC-R2A vs qPCR; HPC-R2A vs FCM-ICC; ATP vs qPCR; ATP vs FCM-ICC; qPCR vs FCM-ICC). Spearman’s correlation coefficients (r) were used for the LRM statistical endorsement. Warning and alarm threshold values, computed by the FDM on single parameters, were applied to each LRM to calculate the corresponding values from linear correlation equations. The mean values (± standard deviation) of four data-driven estimates of warning and alarm thresholds (i.e., one from FDMs plus three from LRMs) of each independent parameter were calculated and presented as alternative standards of water quality. All data elaborations were performed by the software PAST v3.20 (Hammer et al., 2001).

## 3 Results

### 3.1 Heterotrophic plate counts

The number of heterotrophic bacterial colonies (heterotrophic plate count - HPC) varied greatly depending on water origin and across the tested growth conditions, with values ranging from 0 to over 10^6^ CFU/ml. TWs and GWs showed significantly lower values in comparison to RWs and WWs (table 1). Only in TWs, HPC on YEA medium increased significantly passing from 3 to 7 days of incubation (Kruskal-Wallis test, p ≪ 0.01). After the longer incubation time, there was a statistically significant difference between colony numbers found on YEA and R2A and between all water groups (Kruskal-Wallis test, p < 0.05), though HPC from cTWs was close to the method detection limit in all growth conditions. HPC on R2A medium showed values higher than that on YEA medium. FDM thresholds were plotted in figure 1 and reported in table 2. Overall, we found significant differences between contamination levels of all water groups, assessed in terms of cultivability and tested by PERMANOVA (p ≪ 0.001).

**Figure 1.**
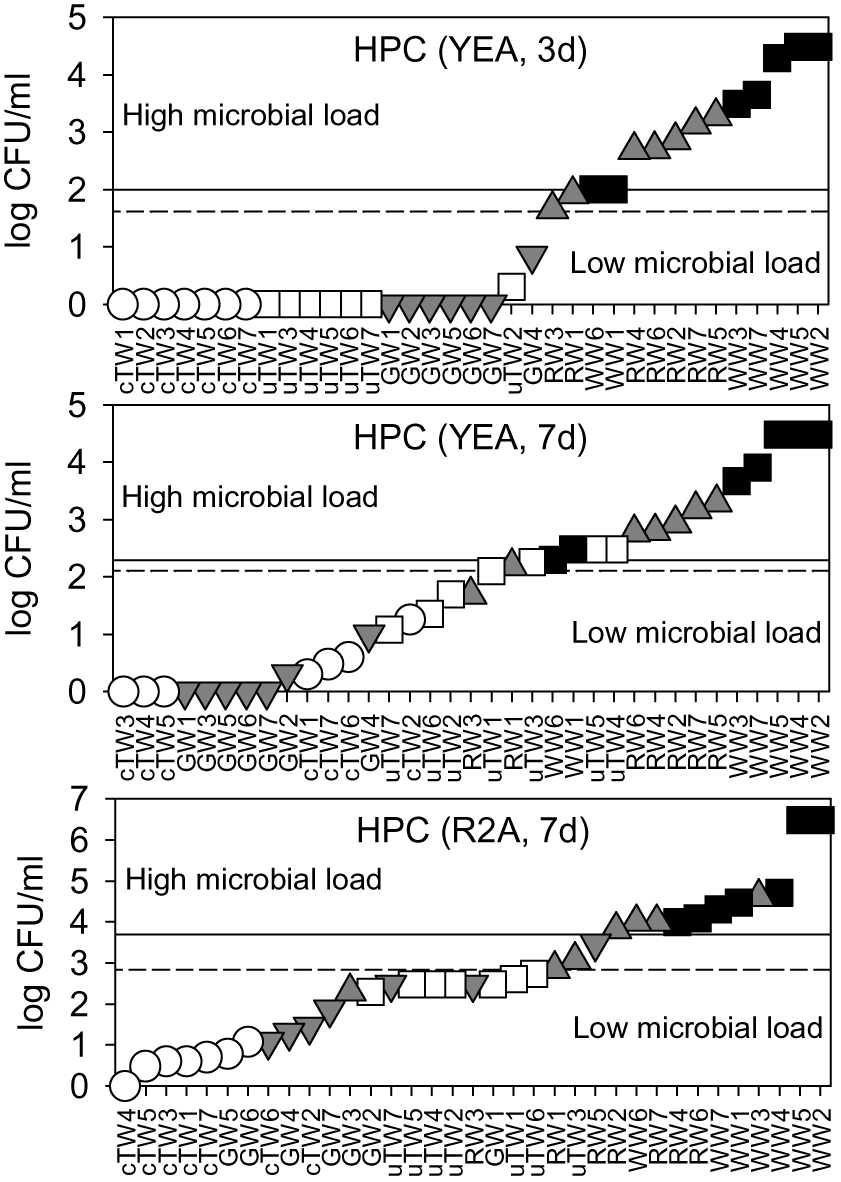
Water microbial load as assessed by plate cultivation on Yeast Agar extract (YEA) and R2A medium, upon 3 days and 7 days of incubation. HPC data were plotted in ascending rank order on a logarithmic scale in order to better visualize warning (dashed lines) and alarm (solid lines) thresholds, which discriminate waters with low and high microbial load. Samples included chlorinated Tap Waters (cTW1-7), unchlorinated Tap Waters (uTW1-7), Ground Waters (GW1-7), Rivers Waters (RW1-7), and Waste Waters (WW1-7).

**Table 1.** Complete dataset of the water microbial load. Heterotrophic Plate Counts (HPC) were assessed from different cultivation media (i.e., YEA, R2A) and incubation times (i.e., 3 and 7 days). Cultivation-independent techniques with current and promising perspectives for space applications were selected (i.e., ATP-metry, qPCR, flow cytometry - FCM). Samples included chlorinated Tap Waters (cTW1-7), unchlorinated Tap Waters (uTW1-7), Ground Waters (GW1-7), Rivers Waters (RW1-7), and Waste Waters (WW1-7).

|  | Cultivation based techniques |  |  | Cultivation independent techniques |  |  |  |
| --- | --- | --- | --- | --- | --- | --- | --- |
|  | HPC YEA-3d<br>CFU/ml | HPC YEA-7d<br>CFU/ml | HPC R2A-7d<br>CFU/ml | ATP-metry<br>pg/ml | qPCR 16S<br>rDNA copies | FCM Total<br>cells/ml | FCM Intact<br>cells/ml |
| cTW1 | 1.00E+00 | 2.00E+00 | 4.00E+00 | 1.48E-03 | 4.21E+02 | 2.55E+04 | 2.23E+04 |
| cTW2 | 1.00E+00 | 1.80E+01 | 2.80E+01 | 2.41E-02 | 1.46E+03 | 4.65E+04 | 2.93E+04 |
| cTW3 | 1.00E+00 | 1.00E+00 | 4.00E+00 | 2.44E-03 | 2.23E+02 | 2.14E+04 | 1.92E+04 |
| cTW4 | 1.00E+00 | 0.00E+00 | 1.00E+00 | 8.02E-03 | 3.70E+02 | 1.50E+04 | 1.30E+04 |
| cTW5 | 0.00E+00 | 0.00E+00 | 3.00E+00 | 3.89E-03 | 5.00E+03 | 5.71E+04 | 5.10E+04 |
| cTW6 | 1.00E+00 | 4.00E+00 | 1.20E+01 | 3.13E-03 | 2.68E+02 | 1.57E+04 | 1.52E+04 |
| cTW7 | 1.00E+00 | 3.00E+00 | 5.00E+00 | 2.46E-03 | 3.70E+02 | 1.79E+04 | 1.55E+04 |
| uTW1 | 0.00E+00 | 1.25E+02 | 4.00E+02 | 1.95E+00 | 1.36E+04 | 1.12E+05 | 8.84E+04 |
| uTW2 | 2.00E+00 | 4.90E+01 | 3.00E+02 | 2.11E+00 | 1.54E+04 | 1.18E+05 | 8.49E+04 |
| uTW3 | 0.00E+00 | 1.83E+02 | 1.20E+03 | 2.75E+00 | 1.76E+04 | 1.45E+05 | 1.03E+05 |
| uTW4 | 1.00E+00 | 3.00E+02 | 3.00E+02 | 2.09E+00 | 1.87E+04 | 1.28E+05 | 1.02E+05 |
| uTW5 | 1.00E+00 | 3.00E+02 | 3.00E+02 | 1.55E+00 | 9.39E+03 | 1.19E+05 | 6.93E+04 |
| uTW6 | 0.00E+00 | 2.30E+01 | 5.40E+02 | 1.41E+00 | 8.85E+03 | 1.25E+05 | 8.94E+04 |
| uTW7 | 0.00E+00 | 1.20E+01 | 3.00E+02 | 1.57E+00 | 1.05E+04 | 1.30E+05 | 8.11E+04 |
| GW1 | 1.00E+00 | 1.00E+00 | 3.00E+02 | 2.18E-01 | 8.64E+03 | 4.61E+04 | 4.37E+04 |
| GW2 | 0.00E+00 | 2.00E+00 | 2.00E+02 | 1.67E-01 | 1.14E+04 | 2.92E+04 | 2.78E+04 |
| GW3 | 0.00E+00 | 0.00E+00 | 2.00E+02 | 7.65E-01 | 1.89E+04 | 6.39E+04 | 6.17E+04 |
| GW4 | 7.00E+00 | 1.00E+01 | 2.00E+01 | 1.95E-01 | 5.69E+03 | 2.48E+04 | 2.37E+04 |
| GW5 | 0.00E+00 | 1.00E+00 | 6.00E+00 | 1.48E+00 | 1.18E+04 | 1.41E+04 | 1.28E+04 |
| GW6 | 0.00E+00 | 0.00E+00 | 1.20E+01 | 2.00E+00 | 1.71E+04 | 1.19E+04 | 1.01E+04 |
| GW7 | 0.00E+00 | 0.00E+00 | 7.30E+01 | 1.43E+01 | 8.88E+04 | 3.42E+04 | 2.92E+04 |
| RW1 | 8.30E+01 | 1.53E+02 | 7.00E+02 | 3.41E+00 | 3.34E+04 | 3.43E+04 | 2.73E+04 |
| RW2 | 7.20E+02 | 8.50E+02 | 6.50E+03 | 2.92E+01 | 1.39E+05 | 1.44E+05 | 1.26E+05 |
| RW3 | 4.50E+01 | 4.90E+01 | 3.00E+02 | 8.66E-01 | 8.67E+03 | 2.09E+04 | 1.93E+04 |
| RW4 | 4.90E+02 | 6.20E+02 | 1.00E+04 | 3.60E+02 | 1.50E+06 | 1.42E+06 | 1.01E+06 |
| RW5 | 1.88E+03 | 1.96E+03 | 2.94E+03 | 3.96E+00 | 2.21E+04 | 3.23E+04 | 2.92E+04 |
| RW6 | 5.00E+02 | 5.90E+02 | 1.20E+04 | 1.58E+02 | 7.49E+05 | 4.32E+05 | 3.84E+05 |
| RW7 | 1.33E+03 | 1.51E+03 | 1.00E+04 | 5.60E+01 | 7.99E+05 | 5.98E+05 | 4.62E+05 |
| WW1 | 1.00E+02 | 3.00E+02 | 3.00E+04 | 1.94E+02 | 2.72E+06 | 2.07E+06 | 1.86E+06 |
| WW2 | 3.00E+04 | 3.00E+04 | 3.00E+06 | 5.90E+03 | 9.81E+07 | 2.92E+07 | 2.56E+07 |
| WW3 | 3.10E+03 | 4.70E+03 | 4.00E+04 | 2.80E+02 | 6.89E+06 | 6.62E+06 | 5.63E+06 |
| WW4 | 1.92E+04 | 3.00E+04 | 5.00E+04 | 2.77E+03 | 7.23E+07 | 1.04E+06 | 9.31E+05 |
| WW5 | 3.00E+04 | 3.00E+04 | 3.00E+06 | 2.72E+03 | 9.15E+06 | 1.51E+07 | 1.20E+07 |
| WW6 | 1.00E+02 | 2.00E+02 | 1.00E+04 | 2.09E+02 | 2.33E+06 | 1.32E+06 | 1.28E+06 |
| WW7 | 4.50E+03 | 7.90E+03 | 2.00E+04 | 3.55E+02 | 3.78E+06 | 2.85E+06 | 2.67E+06 |

### 3.2 Alternative parameters to assess the water microbial load

ATP concentrations varied over more than 6 log units, with a sharp increase from tap waters (range 1.5 - 24.1 x10^-3^ pg ATP/ml) to waste waters (range 1.9 - 59.0 x10^2^ pg ATP/ml) (figure 2a). Apart from TWs and GWs which showed similar values (Kruskal-Wallis test, p > 0.05), the differences in the mean ATP content among water groups were greater than would be expected by chance (Kruskal-Wallis test, p < 0.05).

**Figure 2.**
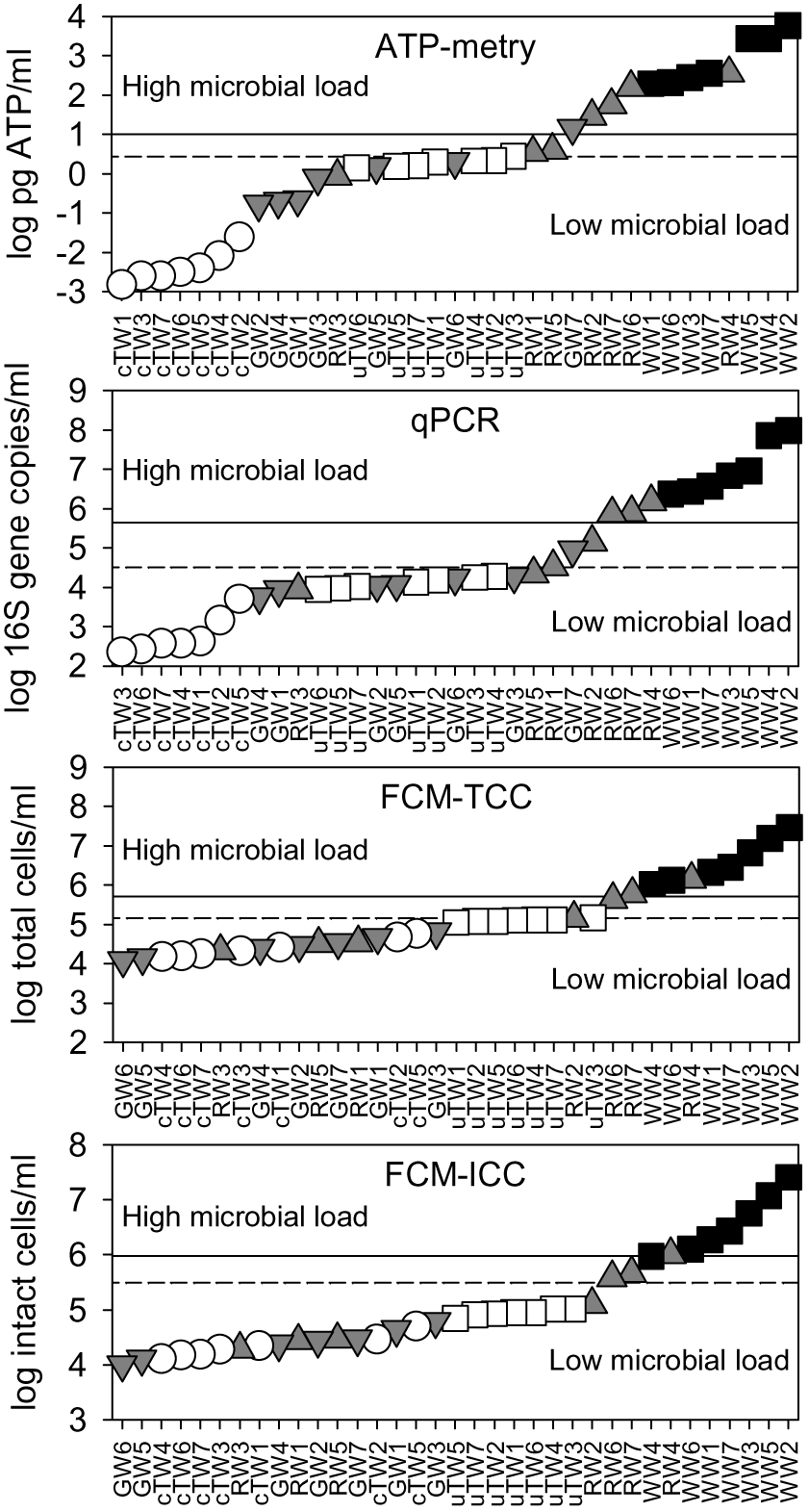
Water microbial load as assessed by alternative parameters (i.e., ATP content, 16S rRNA gene abundance, total cell counts - TCC, intact cell counts - ICC). All data were plotted in ascending rank order on a logarithmic scale in order to better visualize warning (dashed lines) and alarm (solid lines) thresholds, which discriminate waters with low and high microbial load. Samples included chlorinated Tap Waters (cTW1-7), unchlorinated Tap Waters (uTW1-7), Ground Waters (GW1-7), Rivers Waters (RW1-7), and Waste Waters (WW1-7).

As assessed by qPCR, the abundance of 16S rDNA copies varied from 2.2 x10^2^ copies/ml (with minimum values found in chlorinated tap waters) to 9.8 x10^7^ copies/ml (with maximum values found in WWs) (figure 2b). GWs showed similar values to uTWs and RWs (Kruskal-Wallis, p > 0.10), but there were statistically significant differences among all other water groups (Kruskal-Wallis, p < 0.05).

As assessed by flow cytometry, TCC ranged over 3 log units, passing from 1.2 x10^4^ cells/ml (minimum values in GWs) to 2.9 x10^7^ cells/ml (maximum values in WWs) (figure 2c). WWs showed a different mean value from all other groups (Kruskal-Wallis test, p < 0.01), while GWs did not show any statistical difference with cTWs and RWs (Kruskal-Wallis test, p > 0.07). Total cell counts double-checked by epifluorescence microscopy were similar and well correlated with FCM data points on the 1:1 log-log line (Spearman’s r = 0.91, p ≪ 0.001; data not shown). On average, the great majority of total cells comprised membrane-intact cells (ICC = 84.1 ± 10.3 % of TCC), with percentages lower in tap waters (78.1 ± 11.9 %) and higher in ground waters (91.7 ± 5.0 %). Given the limited variation range, ICC followed patterns very similar to TCC on the log scale (figure 2d). FDM thresholds were plotted in figure 2 and reported in table 2. Overall, we found significant differences between the average contamination levels of all water groups, also using the combination of alternative parameters (i.e., ATP, 16S rDNA, ICC) (PERMANOVA, p < 0.01).

**Table 2.** Warning and alarm thresholds, computed according to the frequency distribution model (FDM) of each single parameter (see also figure 1 and 2) and the correlation equations of Linear Regression Model (LRM) between pairs of parameters (see also figure 4). Mean values (± standard deviation) of both thresholds were reported per each parameter and retained as alternative water quality standards.

| Model | Parameter | Warning threshold | Alarm Threshold |
| --- | --- | --- | --- |
| FDM | HPC-YEA3d (CFU/ml) | 41 | 100 |
| FDM | HPC-YEA7d (CFU/ml) | 126 | 195 |
| FDM | HPC-R2A7d (CFU/ml) | 684 | 4898 |
| LRM | HPC vs ATP | 557 | 1604 |
| LRM | HPC vs qPCR | 381 | 4980 |
| LRM | HPC vs FCM | 428 | 4145 |
| | Mean ±sd | 512 ±137 | 3907 ±1580 |
| FDM | ATP (pg/ml) | 2.7 | 10.2 |
| LRM | ATP vs HPC | 3.3 | 26.1 |
| LRM | ATP vs qPCR | 1.7 | 37.9 |
| LRM | ATP vs FCM | 2.0 | 22.2 |
| | Mean ±sd | 2.4 ±0.7 | 24.1 ±11.5 |
| FDM | 16S rDNA ($10^4$ copies/ml) | 3.2 | 44.4 |
| LRM | 16S vs HPC | 5.5 | 29.4 |
| LRM | 16S vs ATP | 4.7 | 13.2 |
| LRM | 16S vs FCM | 3.6 | 29.9 |
| | Mean ±sd | 4.2 ±1.0 | 29.2 ±12.8 |
| FDM | ICC ($10^5$ cells/ml) | 1.0 | 4.2 |
| LRM | ICC vs HPC | 1.3 | 3.6 |
| LRM | ICC vs ATP | 1.2 | 2.1 |
| LRM | ICC vs qPCR | 1.0 | 4.1 |
| | Mean ±sd | 1.1 ±0.2 | 3.5 ±1.0 |

### 3.3 Data correlation and methodological cross-validation

Over the monitored wide range of microbial contamination levels, positive linear correlations were observed between all parameters and methods applied in this study. HPC on R2A showed the best correlations with other parameters in comparison to HPC on YEA. The best fit was found in the log-log linear relation between ATP and HPC-R2A (Spearman’s r = 0.91) and qPCR data (Spearman’s r = 0.97), while the weakest between ATP and FCM data (Spearman’s r = 0.81). HPC-R2A and ICC showed higher correlation coefficients than HPC-YEA and TCC, respectively. Thus, they were used in correlation plots against all other parameters (figure 3).

**Figure 3.**
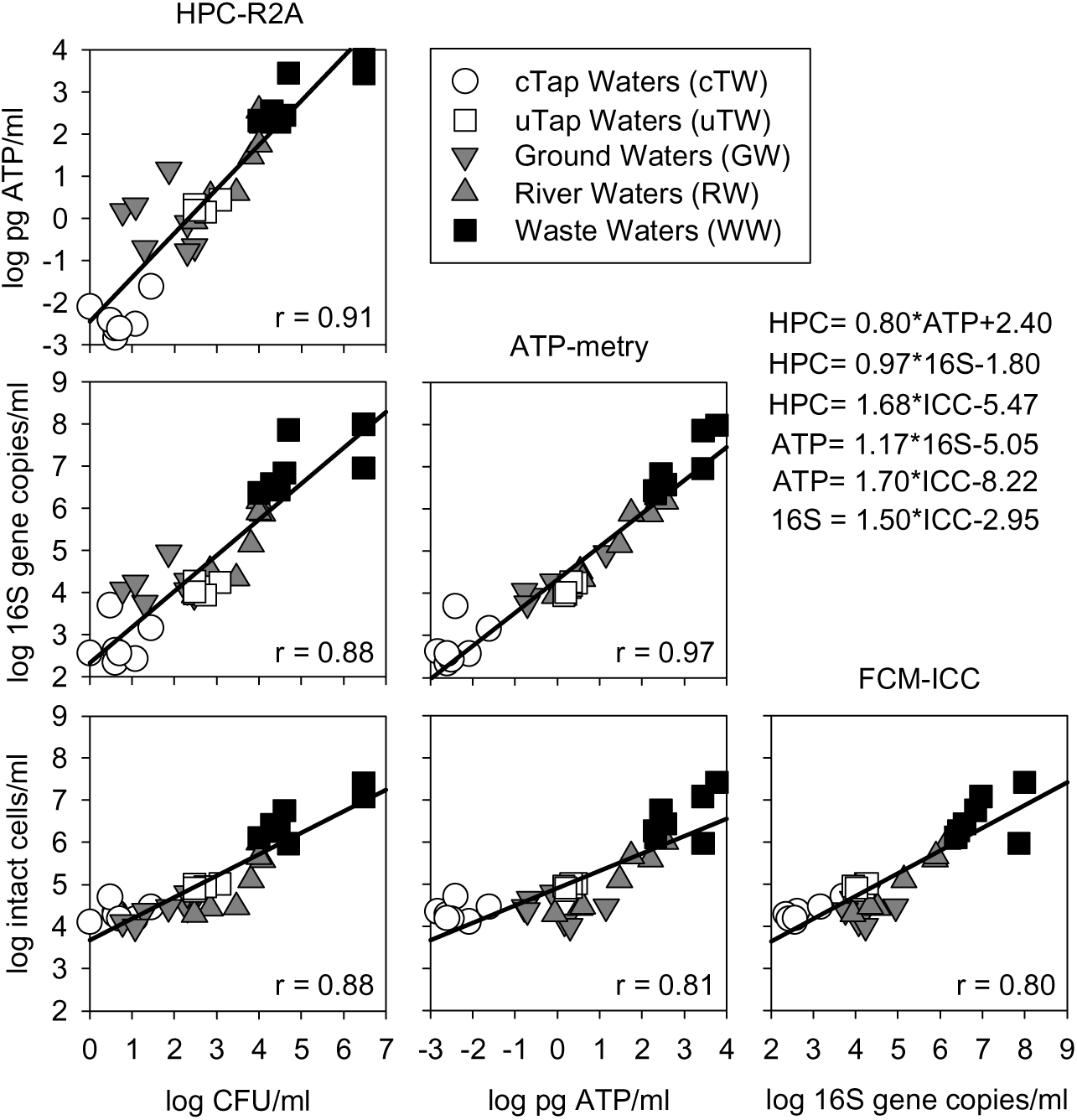
Correlation matrix (log-log) among alternative parameters to assess microbial contamination in waters of different origin (i.e., chlorinated and unchlorinated Tap Waters – cTW and uTW; Ground Waters – GW; River Waters – RW; Waste Waters – WW). Equations of Linear Regression Models (LRMs) are reported, along with Sperman’s correlation coefficients (r).

Warning and alarm thresholds, computed according to the FDM of each single parameter and the LRM correlations equations between all pairs of data (figure 3), were summarized in table 2. The mean values (± standard deviation) of warning and alarm thresholds were proposed as alternative water quality standards based on the microbial load assessed by each independent parameter (table 2).

## 4 Discussion

### 4.1 Suitability of warning and alarm thresholds of water microbial load for space applications

The appropriate consideration of microorganisms in human-confined habitats and their interactions with the space environment are essential to start designing a self-sufficient spacecraft for safe and successful future missions (Pierson, 2001). In extremely confined habitability conditions, as those found onboard crewed space-crafts and during longer-lasting flight missions, water renewal by ground-loaded supplies could be increasingly expensive and unmanageable for months (McCleskey et al., 2012). Therefore, water consumption needs and human health issues may fundamentally rely on a timely detection of unexpected microbiological contamination events, e.g. owing to failure of onboard water recycling and disinfection treatments. Here we presented a data-driven double-threshold procedure intended to identify novel standards of water quality, using alternative cultivation-independent parameters suitable for (near) real-time assessments of the total microbial load.

In current water regulations on Earth, several procedures have been recommended to distinguish between elements of geogenic and anthropogenic origin, elucidate the spatial distribution of chemical elements, identify the source of pollution, and estimate the related risks for human health and activities. The detection of anomalies in the concentration of major and trace chemical elements is one of the main tasks in the adopted statistical approaches (Biddau et al., 2017; Reimann et al., 2005; Stefania et al., 2018). Probability plots, in combination with a data pre-selection, were proposed to graphically represent trends and discontinuities, also identifying data exceeding fixed percentile values (generally the 90^th^, 95^th^, and 97.7^th^ percentile outliers) (Preziosi et al., 2014). In this study, we followed a similar approach but using the microbial load data assessed by different methodologies in order to determine warning and alarm threshold values, respectively set on the 95^th^ and 5^th^ percentiles of values found in the water samples with low and high microbial load. By fundamentally relying on the number and distribution of the available data, the described procedure was not intended to provide fixed limits nor the risks associated with water microbial contamination events. The identified threshold values (table 2) could constitute novel reference values, in view of data deriving from real space conditions and human-confined environments. It is worth noting that two drinking water samples (i.e., Russian potable spring waters), analyzed after 5-years exposure to ISS microgravity conditions by using ATP-metry and flow cytometry with the same full methods herein described, exceeded the alarm thresholds (Bacci et al., 2018 - submitted). Thus, it is likely that some sort of water treatment should be considered to comply with our novel water quality standards.

The microbial load has been retained a key driver of microbial alterations due to varying environmental factors and water treatment settings in numerous studies focused on either natural or engineered aquatic systems (Amalfitano et al., 2018; Besmer and Hammes, 2016; Harry et al., 2016; Osman et al., 2008). However, the application of alternative parameters into regulatory water quality monitoring is still prevented by methodological and procedural issues, including inter-laboratory reproducibility, prioritization of water contaminants, and cross-validation of applied methodologies (Chapman, 1996). In this study, all selected monitoring techniques showed pioneering potential applicability to space and human-confined environments, given the necessity to overcome some basic drawbacks of cultivation-based approaches (i.e., time-to-result up to several days from sampling; growth of opportunistic microorganisms from stored analytical wastes).

### 4.2 HPC acceptability levels in space waters

For preflight and inflight waters, HPC acceptability levels were developed through space analytical experience to mitigate risks to crew health and to maintain the integrity of water treatment systems (e.g., prevention of biofouling in water recirculation and distribution network; microbial growth on hardware components) (Pierson et al., 2013; Van Houdt et al., 2012). According to the ISS Medical Operations Requirements Document (Duncan et al., 2008), ISS waters must be free of coliforms, with a HPC values ≤50 CFU/ml, and sample processing and analysis have to follow precise procedural steps using the US Environmental Health System water kit (NASA, 2005).

When considering the cultivation conditions relatively more similar to those of the US water kit (i.e., HPC on YEA after 3 incubation days at 22°C), TWs and GWs samples accomplished the HPC limit of 50 CFU/ml, with warning and alarm thresholds respectively lower and higher of that established for space requirements. However, the HPC thresholds were considerably higher if estimated from other cultivation conditions (table 2). In drinking water legislation and guidelines, maximum HPC limits can vary from 20 CFU/ml to 500 CFU/ml depending on local regulations and sampling locations (Allen et al., 2004).

In line with recently reviewed data (Diduch et al., 2016), we found that HPC were influenced significantly by cultivation conditions (i.e., HPC-R2A > HPC-YEA), time of incubation (i.e., HPC at 3 days < HPC at 7 days), and the initial microbial load level (figure 1). These results are critical when considering that the ISS is provided with four different supplied waters from the space agencies of USA (NASA), Europe (ESA), Russia (Roscosmos; Russian Federal Space Agency), and Japan (JAXA; Japanese Aerospace Exploration Agency). Water microbial communities (e.g., phylogenetic structure) and local treatment requirements (e.g., addition of different concentration of chlorine, silver, or iodine as biocide agents) may differ considerably, so as HPC outcomes. Further differences in microbial cell cultivability under space conditions are likewise expected owing to microgravity and varying cosmic radiation levels, as it was found either for specific bacterial suspended cultures (Kacena et al., 1999) or for strains isolated from built environments on Earth and cultivated on the ISS (Coil et al., 2016). Experiments have been conducted to improve the speed and efficiency of microbial cultivation assays on the ISS using disposable simple devices and microfluidic systems (e.g., https://www.nasa.gov/mission_pages/station/research/experiments/1033.html; https://www.nasa.gov/mission_pages/station/research/experiments/2357.html). Major advantages arise from target-specific isolation and characterization of different types of microorganisms in pure cultures, including water-borne pathogens (Boitard et al., 2015). However, the HPC reliability for total microbial load assessments in space waters might fall far below the acceptable reproducibility levels, unless other cultivation-independent techniques are applied to provide confirmatory data, as also observed in terrestrial studies.

### 4.3 ATP-metry and advanced automation options for space applications

Based on a 20 years’ experience on space microbial monitoring, ATP-metry has been retained a consistent approach for estimating the viable microbial biomass in water samples (Guarnieri et al., 1997; La Duc et al., 2004). By offering feasible automation options for space applications, we found that ATP-metry allowed to consistently discriminate water types according to their constitutive microbial contamination levels, also showing a wider variation range in comparison to the other selected parameters (figure 2). The highest ratio between alarm and warning thresholds was also observed (table 2).

In drinking water and food industries, routine ATP measurements were added upon commercially available ATP assay kits and compared in-depth to standard cultivation-based outcomes (Bottari et al., 2015; Hammes et al., 2010; van der Wielen and van der Kooij, 2010). One caveat is that community structure variations, with a natural succession of microbial cells with different ATP content (e.g., prokaryotic and eukaryotic cells), may be overlooked owing to the ataxonomic resolution of ATP assays. Therefore, the microbial load evaluations based on ATP-metry could be further strengthened by complimenting with specific cell-targeting parameters (e.g., biomolecular information, total cell counts, cell size measurements) (Siebel et al., 2008; Vang et al., 2014).

### 4.4 Space applicability of qPCR and biomolecular methods

In space research, the successful application of biomolecular assays was found to rely on procedural improvements for extracting cell nucleic acids and selecting appropriate control samples (e.g., with the same amplification efficiency as the target sequence under microgravity conditions), along with instrumental developments (Yamaguchi et al., 2014). In this study, the abundance of 16S rRNA gene copies was significantly different among water types, also showing significant correlations with values of total microbial load assessed by the other parameters (figures 2). The estimated threshold values allowed discriminating waters with low and high microbial load (table 2). Despite showing puzzling low values in TWs and GWs (on average 0.39 ± 0.07 16S rDNA/cell), the 16S rDNA per-cell ratio was highly variable among water types and consistent with literature data (Klappenbach, 2001; Matturro et al., 2013). In view of recent technological developments of molecular methods for space applications, we found that qPCR could be considered as a sensible method for water monitoring, although time-to-results can rise up to several hours from sampling (Lopez-Roldan et al., 2013). Advantages and limitations of the 16S rDNA targeting PCR procedures were reviewed extensively within the context of molecular techniques used to generate data for biomonitoring (Porter and Hajibabaei, 2018; Smith and Osborn, 2009). In particular, it was underlined that current protocols are definitely more informative when used to quantify the occurrence of target functional genes and species of interest (e.g., human pathogens and microorganisms of habitability concerns) rather than estimating the total microbial load (Smith and Osborn, 2009; Sohier et al., 2014).

### 4.5 Flow cytometry: a future alternative tool?

Flow cytometry has been included in the roadmaps of space agencies for monitoring spaceflight-associated requirements (Crucian and Sams, 2005). Though it was not yet specifically tested for onboard water quality assessments, a customized FCM platform was already successfully tested on board the ISS to assess physiological adaptations of astronauts’ blood cells to microgravity (Crucian and Sams, 2012; Dubeau-Laramée et al., 2014; Phipps et al., 2014). In multiple full-scale terrestrial applications, detailed reasons were recognized and meticulously described to argue that FCM could represent a suitable alternative for routine microbiological water monitoring (Van Nevel et al., 2017).

Our results were in line with published data in terms of both total and intact cell counts assessed from different water types including drinking waters (Vital et al., 2012), ground waters (Amalfitano et al., 2014), river waters (Boi et al., 2016), and wastewaters (Foladori et al., 2010). Evidences of significant cross-correlation among microbial quantification techniques are widely reported in literature (Siebel et al., 2008; Vital et al., 2012). Accordingly, we found significant correlations between FCM data and results from epifluorescence microscopy, along with the water microbial load assessed by HPC, ATP-metry, and qPCR (figure 3).

However, unexpected low TCC values particularly in some RWs and WWs could originate from the presence of suspended cell aggregates (i.e., verified by microscopic direct observations), which are acquired as single events (Casentini et al., 2016; Liu et al., 2016). Moreover, the cytometric evaluations are susceptible to increased background levels and debris found in very clean waters (Hammes et al., 2008), with possible TCC over-estimations in TWs. This could partly explain why TCC showed the lower data variation (i.e., 3 log units) in comparison to the other parameters among water types with such different origin and contamination levels. Accordingly, the alarm threshold was only three times higher than the warning threshold. Therefore, FCM routine analysis and developed protocols still require a thorough calibration and validation of their performances and drawbacks for space applications.

## 5 Conclusions

Our results allowed identifying alternative standards of water quality based on the assessment of the water microbial load, thus providing a backbone dataset to develop and test innovative monitoring approaches for space and Earth-based water settings. Cultivation-independent techniques, selected among those consolidated in terrestrial studies and with current or promising perspectives for space applications (i.e., HPC, ATP-metry, qPCR, flow cytometry), could help in managing the ISS on-board water quality, and ultimately enhance the crew responsiveness by providing an immediate alert of microbial load peaks.

## Acknowledgments

This work was supported by the European project BIOWYSE (H2020-COMPET-2015-687447).

